# Draft genome assembly data of *Anoxybacillus sp.* strain MB8 isolated from Tattapani hot springs, India

**DOI:** 10.1101/2021.06.09.447659

**Authors:** P K Vishnu Prasoodanan, Shruti S. Menon, Rituja Saxena, Prashant Waiker, Vineet K. Sharma

## Abstract

Discovery of novel thermophiles has shown promising applications in the field of biotechnology. Due to their thermal stability, they can survive the harsh processes in the industries, which make them important to be characterized and studied. Members of *Anoxybacillus* are alkaline tolerant thermophiles and have been extensively isolated from manure, dairy-processed plants, and geothermal hot springs. This article reports the assembled data of an aerobic bacterium *Anoxybacillus sp.* strain MB8, isolated from the Tattapani hot springs in Central India, where the 16S rRNA gene shares an identity of 97% (99% coverage) with *Anoxybacillus kamchatkensis* strain G10. The de novo assembly and annotation performed on the genome of *Anoxybacillus sp.* strain MB8 comprises of 2,898,780 bp (in 190 contigs) with a GC content of 41.8% and includes 2,976 protein-coding genes,1 rRNA operon, 73 tRNAs, 1 tm-RNA and 10 CRISPR arrays. The predicted protein-coding genes have been classified into 21 eggNOG categories. The KEGG Automated Annotation Server (KAAS) analysis indicated the presence of assimilatory sulfate reduction pathway, nitrate reducing pathway, and genes for glycoside hydrolases (GHs) and glycoside transferase (GTs). GHs and GTs hold widespread applications, in the baking and food industry for bread manufacturing, and in the paper, detergent and cosmetic industry. Hence, *Anoxybacillus sp.* strain MB8 holds the potential to be screened and characterized for such commercially relevant enzymes.

**Specifications Table:** 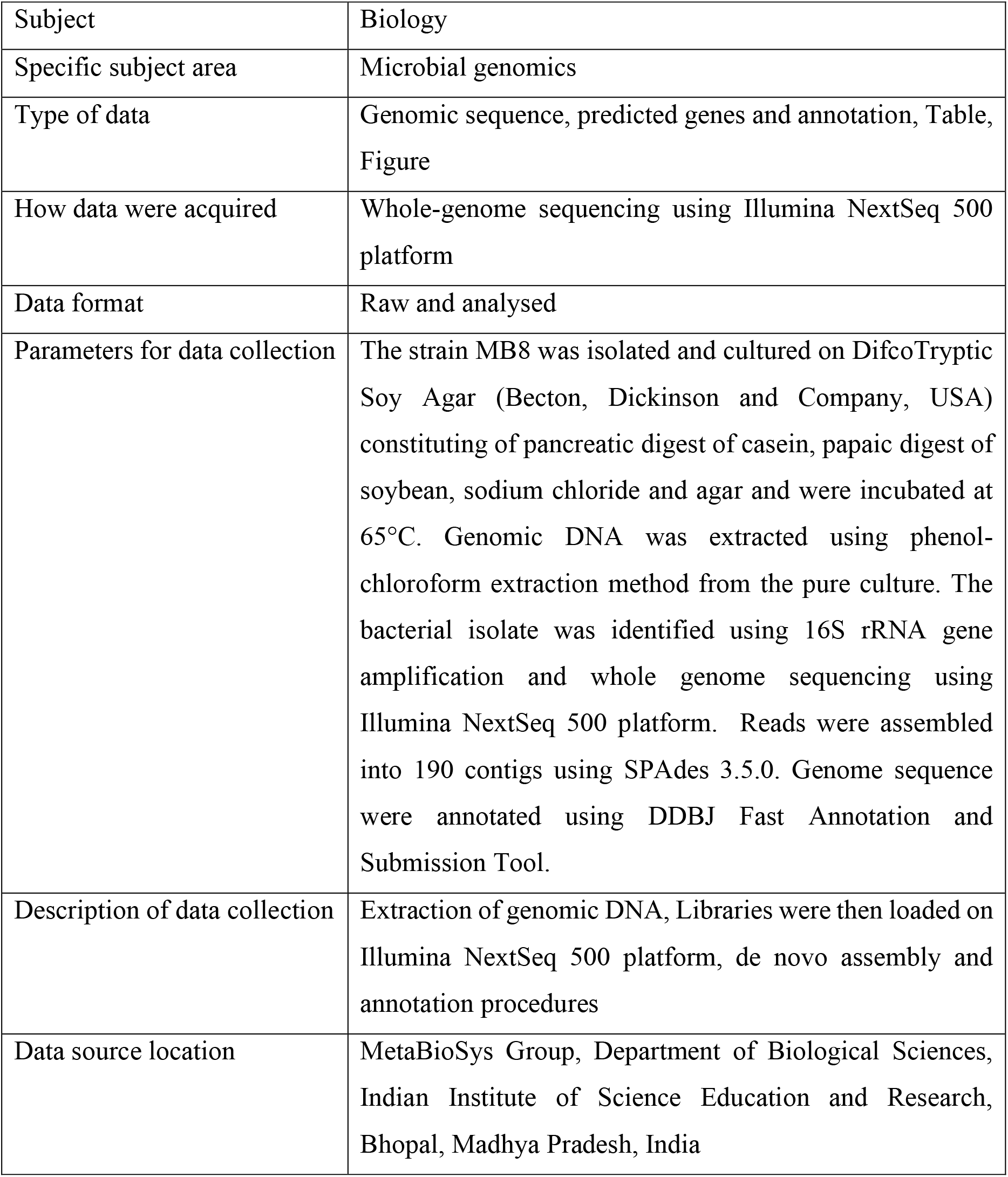

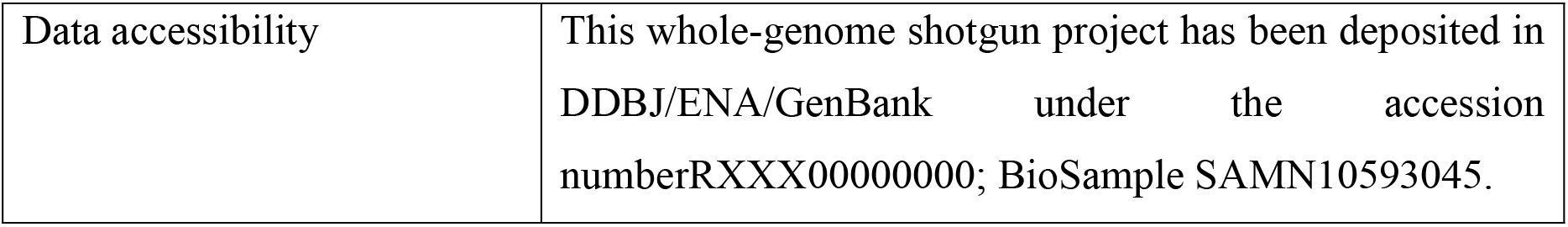

**Values of Data:**

1. The *Anoxybacillus sp.* strain MB8 genome assembly data provides insights into functional potential of thermophilic enzymes of this thermotolerant microbe.
2. The presence of genes for Glycoside Hydrolase (GHs) like alpha-amylase, pullulanase, neopullulanase, alpha-glucosidase, beta-fructofuranosidase etc. and genes for Glycosyl Transferase (GTs) like sucrose synthase, maltodextrin phosphorylase, starch synthase, and glycogen phosphorylase were identified, which hold strong industrial values.
3. The taxonomic annotation of *Anoxybacillus sp.* strain MB8 using different approaches indicates that the closest relatives were *Anoxybacillus gonensis* NZ CP012152T (96.86% ANI) and *Anoxybacillus kamchatkensis* G10 NZ CP025535 (96.83% ANI) obtained from. The strain MB8 most likely belongs to the same subspecies of *Anoxybacillus gonensis* NZ CP012152T.

## 1. Data Description

Thermophiles are known for their capability to subsist in extreme temperatures and are known to be relevant in industrial and biotechnological applications. Metagenomic analysis of several extreme ecosystems across the globe has revealed micro-organisms harbouring genes for enzymes showing a promising role in the applied biotechnology [1–3]. India is known to harbour about 400 thermal springs according to the Geological Survey of India [4], and a study conducted at Tattapani hot-springs (located in the state of Chhattisgarh, India) revealed to harbor bacterial species that have adapted to survive in extreme ecosystems [1]. *Anoxybacillus mongoliensis* strain MB4 was previously isolated from the same hot spring and is a sulfur-utilizing aerobic thermophile [5]. The genome sequence of Gulbenkiania mobilis and Tepidimonas taiwanensis, chemolithotrophic thermophiles, have been reported previously from the Anhoni hot springs located in Central India [6,7]. Here, we report the draft genome of *Anoxybacillus sp.* strain MB8, isolated from Tattapani hot springs with a temperature of 61.5°C, located in the state of Chhattisgarh, Central India (23.41°N latitude and 83.39°E longitude). The 16S rRNA gene sequence of MB8 showed 97% sequence identity with the thermophilic bacteria A. kamchatkensis strain G10, previously isolated from a hot spring in Indonesia [8]. The genome shows the presence of assimilatory sulfate reduction pathway and the presence of glycoside hydrolases (GHs) and glycosyltransferases (GTs). The GH enzymes are capable of degrading carbohydrates and starch; which are the major raw materials in industries. Glycoside hydrolases have been widely used in food manufacturing, cosmetics industry, and animal nutrition. These enzymes have been predicted to have important roles in biorefining applications, food, and baking industries as well as in second-generation biofuel production [9–11].

Previous studies have reported, *Anoxybacillus spp.* to possess genes for starch modifying enzyme. The thermostable properties of the enzymes make them a suitable and stable catalyst for industrial applications, and thus their study and characterization seem to be important for their efficient utilization. This article also reports reference-based genome alignment of the strain MB8 with the five complete and twenty-five available draft genome sequences of the members of genus *Anoxybacillus*. The analysis reveals the alignment of *Anoxybacillus sp.* MB8 with *Anoxybacillus kamchatkensis* strain G10 (73.77%) to be the highest among the known five complete genomes whereas *Anoxybacillus gonensis* strain DT3-1 (72.25%) to be highest among the draft genomes. The MiGA, digital DNA: DNA hybridization (dDDH) obtained using genome-to-genome distance calculator (GGDC) and average nucleotide identity (ANI) revealed that the *Anoxybacillus sp.* MB8 most likely is a subspecies of *Anoxybacillus gonensis* NZ CP012152T.

The colonies grown on DifcoTryptic Soy Agar were medium-sized, round-shaped, and transparent with irregular margins. The growth of the colonies was observed after 12-16h of incubation under aerobic conditions. The 16S rDNA sequence was used to identify the species. The isolate showed a 97% sequence identity and 99% coverage with *Anoxybacillus kamchatkensis* strain G10. The phylogenetic tree (**Fig. 1**) was used to reveal its position in the taxonomic tree, which confirmed that strain MB8 is a member of the genus *Anoxybacillus*.

**Fig. 1:**
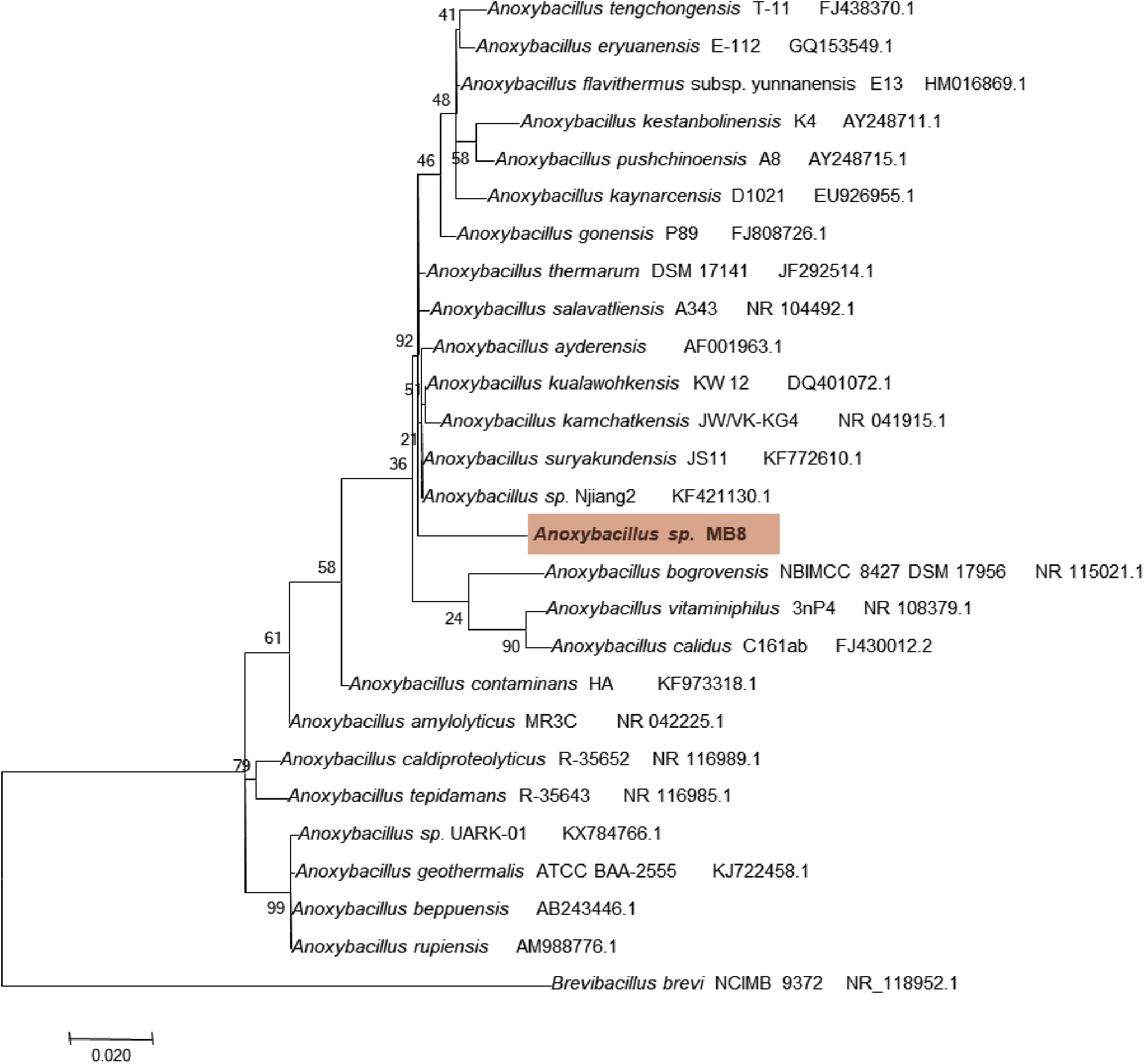
Phylogenetic analysis of *Anoxybacillus sp.* strain MB8 using the Maximum Likelihood method. The tree was constructed based on the 16S rRNA gene sequence. The scale bar represents 0.02 nucleotide substitutions per position and the species used as an out-group is *Brevibacillus brevis* NCIMB 9372. The bootstrap consensus tree inferred from 1000 replicates and the branches corresponding to partitions reproduced in less than 50% bootstrap replicates are collapsed. The evolutionary analyses were conducted in MEGA7.

Illumina sequencing generated 849.3 MB of total data and 47, 86,534 reads (trimmed and filtered) were used for the genome assembly using SPAdes 3.5.0. The draft genome sequence annotated by DFAST server contains 190 contigs (length>=300bp, k-mer set to 91) with an N50 of 54,614 bp and 41.8% G+C content. A total of 2,976 protein-coding genes, RNA genes (1 rRNA operon, 73 tRNA, and 1tmRNA) and 10 CRISPR array were predicted (**Table 1**). A total of 2,767 protein coding-genes were assigned to 21 eggNOG categories with their relative abundance after alignment of the genes with eggnog v4.1 database and assigning eggNOG IDs. Within the 21 categories, the majority of genes were assigned to “Function unknown” (26.3823%) followed by “General function prediction only” (7.58%) and “Replication, recombination and repair” (7.37%). The least number of genes were assigned to “Chromatin structure and dynamics” (0.018%). The relative abundance of each category is shown in (**Table 2**).

**Table 1:** Genome Features of *Anoxybacillus sp.* MB8

| Attribute | Value |
| --- | --- |
| genome size | 2,898,780 bp |
| Total no. of contigs | 190 contigs (length>=300) |
| Shortest contig | 300 |
| Longest contig | 1,60,603 |
| N50 | 54,614 bp |
| average length | 15,257 |
| GC content | 41.8% |
| Total no. of protein-coding genes | 2,976 |
| Total no. of rRNA operon | 1 |
| Total no. of tRNA | 73 |
| Total no. of tmRNA | 1 |
| Total no. of CRISPRS | 10 |
| Coding Ratio | 88.1% |

**Table 2:** eggNOG functional categories obtained using the eggNOG database

| Code | Relative abundance (%) | Description |
| --- | --- | --- |
| B | 0.0180 | Chromatin structure and dynamics |
| C | 5.5836 | Energy production and conversion |
| D | 1.0842 | Cell cycle control, cell division, chromosome partitioning |
| E | 7.0654 | Amino acid transport and metabolism |
| F | 2.3671 | Nucleotide transport and metabolism |
| G | 3.6863 | Carbohydrate transport and metabolism |
| H | 3.6320 | Coenzyme transport and metabolism |
| I | 3.0538 | Lipid transport and metabolism |
| J | 5.3306 | Translation, ribosomal structure, and biogenesis |
| K | 5.7824 | Transcription |
| L | 7.3726 | Replication, recombination and repair |
| M | 3.8670 | Cell wall/membrane/envelope biogenesis |
| N | 1.4395 | Cell Motility |
| O | 3.4815 | Posttranslational modification, protein turnover, chaperones |
| P | 5.3668 | Inorganic ion transport and metabolism |
| Q | 0.7228 | Secondary metabolites biosynthesis, transport, and catabolism |
| R | 7.5894 | General function prediction only |
| S | 26.3823 | Function unknown |
| T | 3.9573 | Signal transduction mechanisms |
| U | 0.8794 | Intracellular trafficking, secretion, and vesicular transport |
| V | 1.3371 | Defense mechanisms |

As revealed by the KAAS analysis, the genome MB8 contained genes coding for a complete set of enzymes of general carbohydrate metabolism namely, glycolysis, the TCA cycle, and the pentose phosphate pathway. The genes for catalase-peroxidase, thiol peroxidase, glutathione peroxidase and (Fe-Mn, Cu-Zn) superoxide dismutase was also observed, which are important for protection against reactive oxygen species. Genes for menaquinol-cytochrome c oxidoreductase (qcrA, qcrB, qcrC) were identified, which seem to be absent in A. kamchatkensis strain G10 [8,12]. Precursors for carotenoid biosynthesis are formed by the MEP/DOXP pathway. The only carotenoid biosynthesis genes present were crtB, crtN, and crtP, which can be correlated to the colorless appearance of the colonies. Genes for sugar phosphotransferase systems (PTS) like glucose, sucrose, mannitol, and trehalose were found, indicating the ability of the bacteria to utilize multiple carbohydrates as a sole source of carbon. The genes for assimilatory sulfate reduction pathway (sulfate to sulfide conversion), dissimilatory nitrate reduction to ammonium (NarGHI, NrfAH), base excision repair, and nucleotide excision repair were also observed. The presence of dissimilatory nitrate reduction indicates that the bacterium utilizes nitrate or nitrite as an electron acceptor during anaerobic conditions [13].

Genes involved in the selenocompound metabolism like PAPSS and TXNRD, which help in the conversion of selenate to hydrogen selenide were identified. Also, genes involved in the conversion of selenocysteine to seleno-methionine were present. The genome comprised of genes kinA, kinB, kinD, kinE, spo0B, spo0Fand spo0A (a specific master regulator gene) present in sporulation biofilm formation [14]. Genes for aerobic and anaerobic respiration in case of oxygen limitation and genes involved in flagellar motor switch adaption were identified. The genome showed the presence of genes for GHs like alpha-amylase, pullulanase, neopullulanase, alpha-glucosidase, beta-fructofuranosidase, 6-phospho-β-glucosidase, cyclomaltodextrinase (CDase), 1, 4-alpha-glucan branching enzyme, trehalose-6-phosphate hydrolase and oligo-1, 6-glucosidase, which hold strong industrial value [15]. Genes for GTs like sucrose synthase, maltodextrin phosphorylase, starch synthase, and glycogen phosphorylase were identified. There were genes encoding few prokaryotic type-ABC transporters like arabinooligosacchrides, ribose, zinc, etc. Presence of phosphotransacetylase, acetate kinase, and L-lactate dehydrogenase genes corresponds to fermentative growth in B. subtilis and A. flavithermusWK1[12], all of which were identified in the strain MB8.

Reference-based genome alignment for *Anoxybacillus sp.* strain MB8 was carried out using software Bowtie2 [16] and BWA [17]. Trimmed reads (47,86,534) were aligned against complete genomes of five *Anoxybacillus spp.* as well as the twenty-five draft genomes of *Anoxybacillus spp.* The alignment rate against the five complete genomes was obtained using the Bowtie2 (**Table 3, Fig. 2a**) and the MB8 genome aligned highest with *Anoxybacillus kamchatkensis* strain G10 (73.77%) and the lowest with *Anoxybacillus sp.* strain B7M1 (2.84%). The reads were also aligned with the 190 contigs obtained after assembly, which showed a 98.68% alignment rate. The alignment rate against the 25 draft reference genomes varied from 1.25%-72.25% (**Fig. 2b**). The alignments obtained from BWA (**Fig. 3 and 4**) were further analyzed by Qualimap [18]. The comparison between the 5 reference genomes clearly reveals the coverage (X) across the genome is higher for A. kamchatkensis strain G10 and lowest for *Anoxybacillus sp*. strain B7M1. The coverage histograms (**Fig. 4**) indicate the number of genomic locations vs. coverage (X). The comparison between the complete genomes indicates *A. kamchatkensis* strain G10 having better coverage (X= 208-271) for number base pairs, whereas *A. amylolyticus* strain DSM, *Anoxybacillus sp.* strain B7M1, *A. flavithermus* strain WK1 have lower coverage. *A. gonensis* strain G2 also has a similar coverage histogram as that of *A. kamchatkensis* strain G10.

**Table 3:** Alignment rate of reads against complete genomes of *Anoxybacillus* available in NCBI using Bowtie2

| Genome name | Alignment rate | Total aligned reads |
| --- | --- | --- |
| *Anoxybacillus* sp. strain B7M1 | 2.84% | 1,36,014 |
| *Anoxybacillus amylolyticus* strain DSM | 4.53% | 2,16,934 |
| *Anoxybacillus flavithermus* strain WK1 | 24.60% | 11,77,579 |
| *Anoxybacillus gonensis* strain G2 | 73.54% | 35,20,021 |
| *Anoxybacillus kamchatkensis* strain G10 | 73.77% | 35,31,006 |
| *Anoxybacillus* sp. strain MB8 | 98.68% | 47,23,548 |
|  | Total no. of reads | 47,86,534 |

**Fig. 2:**
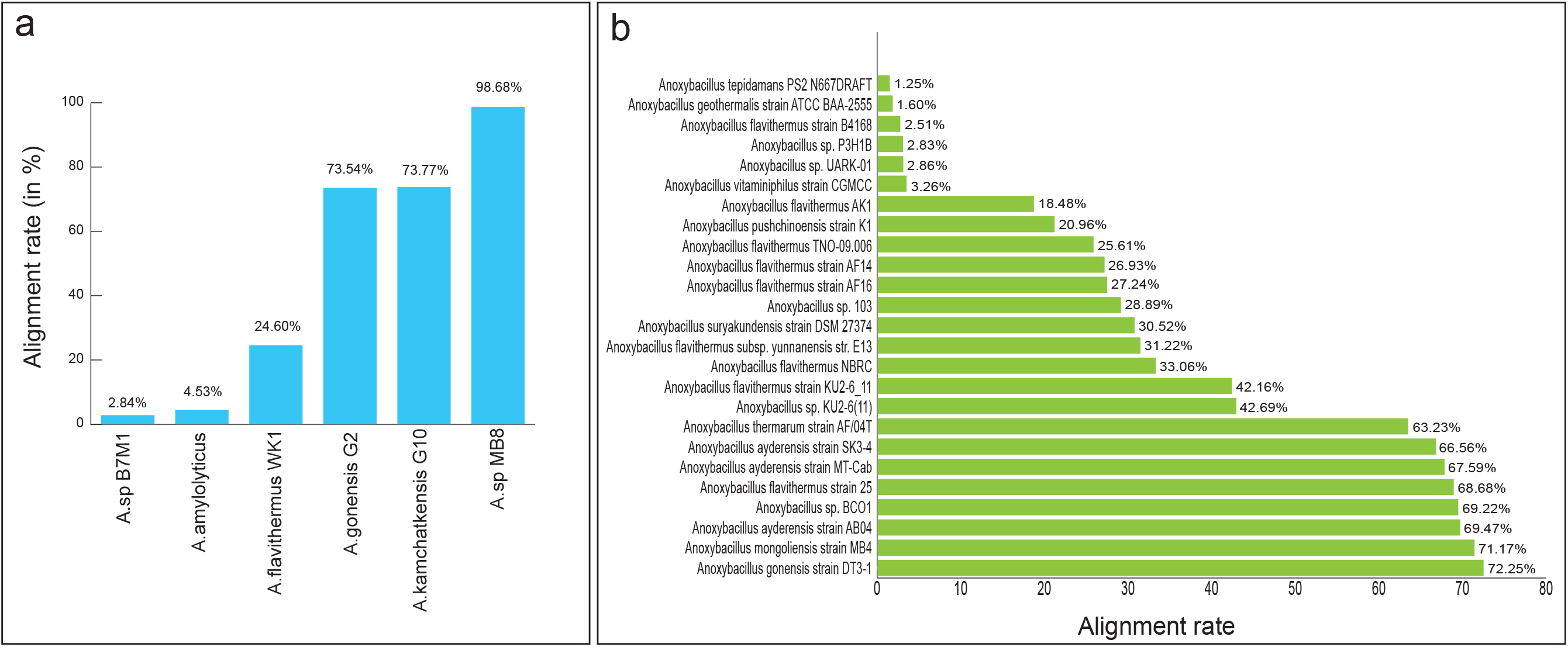
Alignment of *Anoxybacillus sp.* strain MB8 reads against complete and draft genomes of *Anoxybacillus* genus. The software Bowtie2 was used to calculate the alignment rates (in percentage). a) The column chart explains the alignment rate of the *Anoxybacillus sp.* strain MB8 aligned against complete genomes of *Anoxybacillus sp.* strain B7M1, *Anoxybacillus amylolyticus* strain DSM, *Anoxybacillus flavithermus* strain WK1, *Anoxybacillus gonensis* strain G2, *Anoxybacillus kamchatkensis* strain G10, and *Anoxybacillus sp.* strain MB8. b) The bar chart depicts the alignment rate of the *Anoxybacillus sp.* strain MB8 aligned against twenty-five draft genomes of *Anoxybacillus* species downloaded from NCBI

**Fig. 3:**
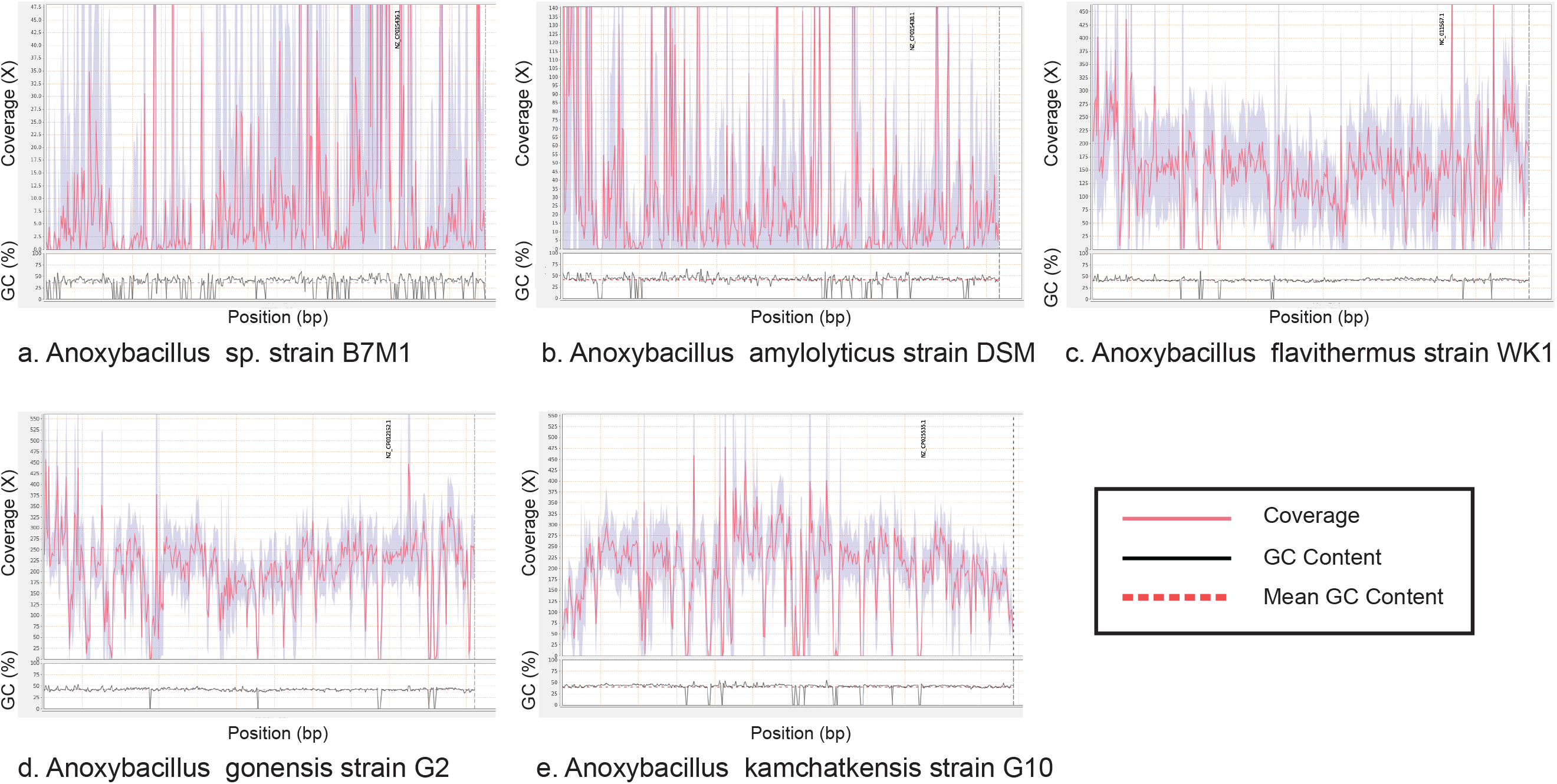
Coverage across reference genomes using Qualimap. It represents the coverage distribution and deviation across the reference. The bottom part of each figure indicates the GC content across a reference (black line) and the dotted line shows its average value. The genome *Anoxybacillus sp.* strain MB8 was aligned against the 5 complete genomes using Burrows-Wheeler Aligner (BWA).

**Fig. 4:**
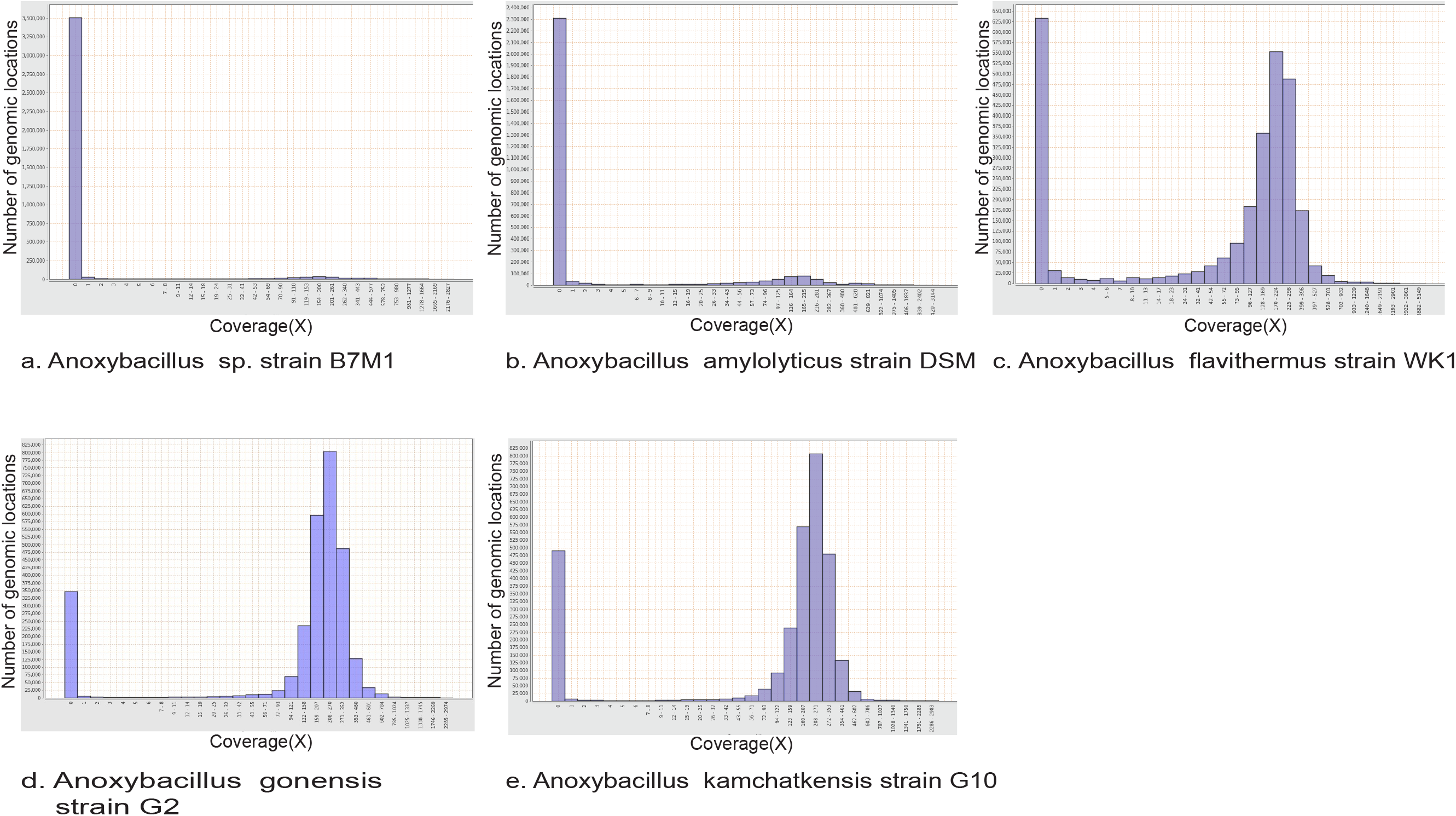
Genome coverage histogram generated from Qualimap. This figure is created after aligning genome MB8 reads with the 5 reference genomes using BWA.

According to the 95% ANI and 70% dDDH, which are the thresholds for species demarcation [19], the OGRI (dDDH and ANI) results revealed that the *Anoxybacillus spp.* strains MB8, G2 and G10 have a high degree of genomic relatedness. The results indicate that *Anoxybacillus gonensis* G2 shares 71.1% dDDH and 96.73% ANI whereas *Anoxybacillus kamchatkensis* G10 shares 71.8% dDDH and 96.67% ANI with *Anoxybacillus sp.* strain MB8.

The MiGA results also indicate that the closest relatives were *Anoxybacillus gonensis* NZ CP012152T (96.86% ANI) and *Anoxybacillus kamchatkensis* G10 NZ CP025535 (96.83% ANI) obtained from MiGA. The strain MB8 most likely belongs to the same subspecies of *Anoxybacillus gonensis* NZ CP012152T (**Fig. 5**).

**Fig. 5:**
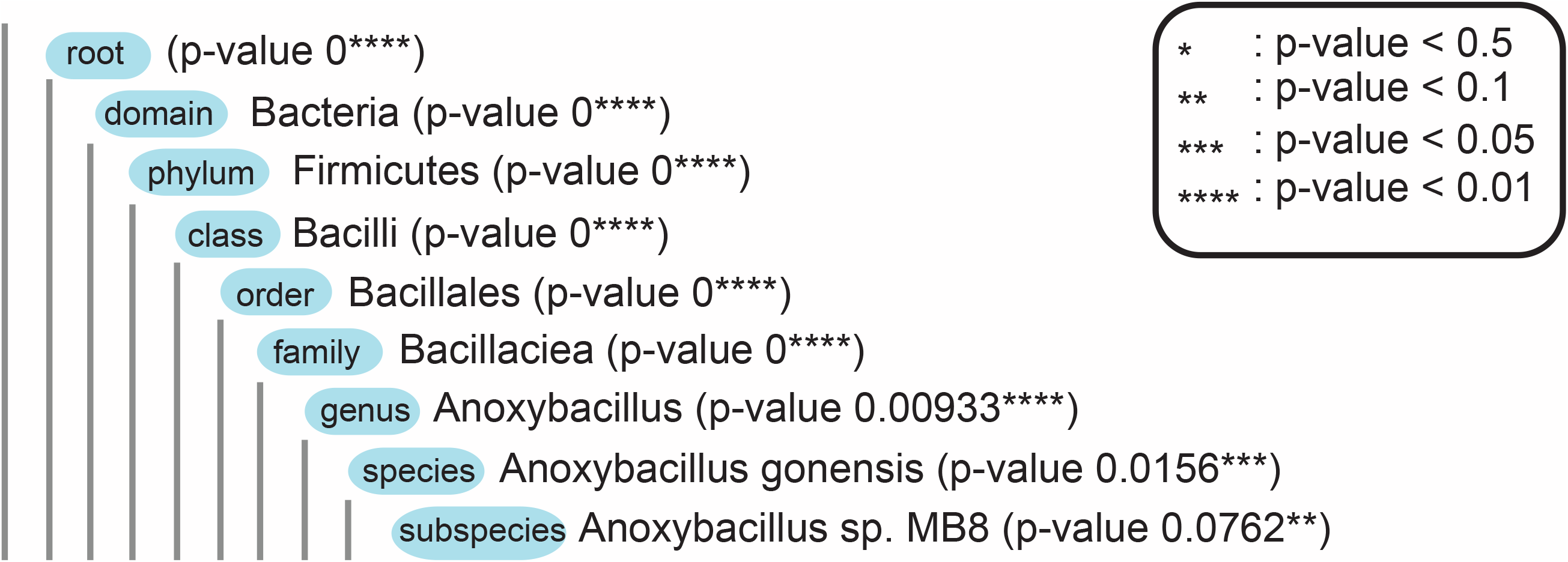
Figure showing the taxonomic classification of reads using Microbial Genomes Atlas (MiGA) online service. It represents that the isolate MB8 is closely related and most likely be a subspecies of Anoxybacillus gonensis.

## 2. Experimental Design, Materials and Methods

### 2.1. Growth conditions

The strain MB8 was isolated and cultured on DifcoTryptic Soy Agar (Becton, Dickinson and Company, USA) constituting of pancreatic digest of casein, papaic digest of soybean, sodium chloride and agar and were incubated at 65°C.

### 2.2. DNA extraction, PCR amplification, and Phylogenetic analysis

Genomic DNA was extracted using phenol-chloroform extraction method from the pure culture. The bacterial isolate was identified using 16S rRNA gene amplification and sequencing using universal eubacteria specific primers (8F: 5’-AGAGTTTGATCCTGGCTCAG-3’ and 1492R: 5’-GGTTACCTTGTTACGACTT-3’) [20]. The PCR product was analyzed by gel electrophoresis, which was observed to be of ~1400 bp size. The amplified regions were sequenced using Sanger sequencing by ABI 3500 genetic analyzer sequencer (Applied Biosystems) at the DNA sequencing facility of IISER Bhopal. The forward and reverse sequences obtained were assembled using FLASH [21]. A similarity search against the NCBI database was performed for each of the assembled sequences using BLASTn [14]. Further, the phylogenetic analysis was carried using the MEGA7 software [22].

### 2.3. Genome sequencing, de novo assembly, and annotations

The genomic libraries were prepared using the Illumina Nextera XT sample preparation kit (Illumina Inc., USA). The libraries were evaluated on 2100 Bioanalyzer using Bioanalyzer High Sensitivity DNA kit (Agilent Technologies, Santa Clara, CA, USA) to estimate the library size. They were further quantified on a Qubit 2.0 fluorometer using Qubit dsDNA HS kit (Life technologies, USA) and by qPCR using KAPA SYBR FAST qPCR Master Mix and Illumina standards and primer premix (KAPA Biosystems, Wilmington, MA, USA) following the Illumina suggested protocol. Libraries were then loaded on Illumina NextSeq 500 platform using NextSeq 500/550 v2 sequencing reagent kit (Illumina Inc., USA) and 150 bp paired-end sequencing was performed at the Next-Generation Sequencing (NGS) Facility, IISER Bhopal, India.

The pre-processing, i.e., ambiguity filtration and quality control, of the raw reads obtained from Illumina sequencing, was performed using the NGSQC Toolkit_v2.3.3 [23]. The filtered reads obtained with a cut-off value for read-length 70 and PHRED quality score of 25 were selected for de novo assembly, which was carried out using SPAdes 3.5.0 [24]. The k-mer parameters from 21 to 101 were used to assemble the reads. The contigs assembled with the k-mer 91 were selected for gene prediction and annotation. The genome sequences (190 contigs) were annotated using DDBJ Fast Annotation and Submission Tool (DFAST) [25]. The tRNA genes were predicted using tRNAscan-SE 2.0 web server [26]. The KEGG Automated Annotation Server (KAAS) analysis was used to assign KOs and to identify metabolic pathways [27]. Further, the annotated genome with 2,796 genes was assigned eggNOG categories by aligning them against eggnog v4.1 database [28] using NCBI-blast-2.6.0+ [29] and were annotated with eggNOG ID. The representation of eggnog classes was deducted using in-house Perl scripts. Contigs were further assembled (72 scaffolds) using Medusa web server [30] for predicting rRNA genes using RNAmmer 1.2 Server [31].

### 2.4. Alignment of MB8 reads against complete genome of Anoxybacillus spp

The complete genome sequences of *Anoxybacillus sp.* strain B7M1, *Anoxybacillus amylolyticus* strain DSM, *Anoxybacillus flavithermus* strain WK1, *Anoxybacillus gonensis* strain G2, and *Anoxybacillus kamchatkensis* strain G10 as well as the twenty-five available draft genome sequences were retrieved from NCBI database. Reference-based alignment was carried out using the reads of genome MB8 with the tool Bowtie2 and Burrows-Wheeler Aligner [16,17]. Reads were also aligned against the 190 contigs (length>=300bp) of *Anoxybacillus sp.* strain MB8. Qualimap [18] was used for comparing the alignment coverage and GC content after aligning the reads against an aforementioned complete reference genomes. To determine the closest genomic neighbours and nucleotide identities, the web server Microbial Genomes Atlas (MiGA) was used, where the query genome MB8 was searched against the NCBI prokaryotic genome database [32]. The Overall Genome Relatedness Indices (OGRI) was calculated between the query and the 5 reference genomes using Genome-to-Genome Distance Calculator (GGDC) version 2.1 online server, using the recommended formula 2 [33]. The OrthoANIu tool [34] was used to calculate the Average Nucleotide Identity (ANI) values between the query MB8 and two complete reference genome i.e. *Anoxybacillus gonensis* G2 and *Anoxybacillus kamchatkensis* G10. The 190 contigs (>=300bp) obtained after assembly was used for MiGA, ANI and GGDC calculations.

## Author Contributions

RS, PW, and VKS carried out the sample collection. RS and PW cultured and isolated the strain, and carried out the 16S rRNA sequencing and identification. RS prepared the sequencing libraries and carried out the whole genome sequencing. VPPK and SSM estimated the genome size and assembly, assessment of assembly quality, carried out the genome annotation, eggNOG analysis, and reference-based genome analysis. SSM carried out the MiGA, GGDC, and OrthoANIu analysis. SSM, RS, VPPK, and VKS wrote the manuscript. All authors read, edited, and approved the final manuscript.

## Acknowledgments

The authors gratefully acknowledge the NGS facility at IISER Bhopal for providing the infrastructure to carry out the sequencing experiments. RS, SSM and VPPK acknowledge DST-INSPIRE for providing research fellowship. We thank MHRD, Govt of India for funding Centre for Research on Environment and Sustainable Technologies (CREST) at IISER Bhopal for providing financial support. However, the views expressed in this manuscript are that of the authors alone and no approval of the same, explicit or implicit, by MHRD should be assumed.

